# Descriptron-GBIF Annotator: A browser-based platform for crowdsourced morphological annotation of biodiversity images to help accelerate morphology based biodiversity data

**DOI:** 10.64898/2026.03.10.710887

**Authors:** Alex R. Van Dam, Francisco Hita Garcia

## Abstract

The accelerating biodiversity crisis demands new approaches to taxonomic description that can scale beyond the capacity of professional taxonomists alone. We present the Descriptron-GBIF Annotator, a zero-installation, browser-based tool for morphological annotation of biodiversity specimen images retrieved directly from the Global Biodiversity Information Facility (GBIF). The application runs entirely client-side as a single HTML file, integrating SAM2 (Segment Anything Model 2) for AI-assisted segmentation, ontology-linked anatomical region templates covering 25 major taxonomic groups across 124 standardized views, 335 ontology Compact URI Expressions (CURIEs), with 745 possible ontology mentions, and structured trait attribute recording. The annotator supports multiple export formats including Darwin Core JSON, COCO JSON, traits CSV, and a novel JSON-LD knowledge graph linking specimens to anatomical regions and morphological traits via UBERON and domain-specific ontologies. A built-in Zenodo publishing pipeline enables users to deposit annotations as citable datasets with DOIs directly from the browser. Additionally users can also annotate images from Zenodo BioSysLit enabling annotation of taxonomic treatments directly. We position this public-facing tool as the first tier of a two-tier architecture complementing the Descriptron Portal, a GPU-accelerated professional workbench for taxonomists providing tools for fine-tuning AI models, geometric morphometrics, and automated species descriptions. Together, these tiers create a feedback loop where public annotations generate training data for expert AI models, while expert-validated outputs improve the public tool. This approach draws on the citizen science model pioneered by Notes from Nature and iNaturalist to engage diverse audiences in structured morphological data collection, addressing a critical gap in biodiversity informatics where specimen images exist in abundance but structured morphological annotations remain scarce. To learn more go here: https://descriptrongbifannotator.org

## Introduction

There is a growing gap between the number of species requiring description and the number of trained taxonomists available to describe them, which represents one of the most pressing challenges in biodiversity science (Engel et al., 2021; Wheeler et al., 2004). Conservative estimates suggest that 80% of the world’s species remain undescribed (Mora et al., 2011), with the situation most acute among hyperdiverse arthropod families sometimes termed *dark taxa* where entire clades containing thousands of species lack any specialist (Hartop et al., 2022). The Global Biodiversity Information Facility (GBIF) has mobilized over 2.7 billion occurrence records, yet for most specimens the available data consists primarily of locality, date, and taxonomic assignment with no structured morphological information (GBIF, 2026, 2011).

Simultaneously, citizen science platforms have demonstrated remarkable capacity to engage public participants in biodiversity data collection. iNaturalist has accumulated over 200 million observations with community-based identifications (iNaturalist, 2026; Van Horn et al., 2018), while Notes from Nature pioneered the crowdsourcing of museum specimen label transcription (Hill et al., 2012). However, these platforms focus primarily on occurrence data and label transcription rather than structured morphological annotation. The result is a paradox: millions of high-resolution specimen images are publicly accessible through GBIF and institutional repositories, yet no widely-available tool enables non-specialists to contribute structured morphological observations about what they see in these images.

Computer vision has advanced rapidly in recent years, with foundation models such as the Segment Anything Model (SAM) and its successor SAM2 (Ravi et al., 2024) achieving remarkable zero-shot segmentation across domains including biological specimens. The DINOv2/v3 family of self-supervised vision transformers (Oquab et al., 2023; Siméoni et al., 2025) has demonstrated powerful feature extraction applicable to fine-grained visual recognition tasks. These models, combined with efficient deployment via ONNX runtime web infrastructure could make it possible for biodiversity science to take advantage of their utility in segmenting biological images.

We present the Descriptron-GBIF Annotator, a comprehensive browser-based tool that bridges this gap by enabling anyone with a web browser to create structured, ontology-linked morphological annotations of GBIF specimen images with AI assistance. We describe its design as the public-facing first tier of a two-tier system complementing the Descriptron Portal, a professional-grade morphological analysis workbench, and discuss its potential utility as a contribution to biodiversity informatics.

## Methods

### System description

#### Architecture and deployment

The Descriptron-GBIF Annotator is implemented as a single, self-contained HTML file (approximately 270 KB) containing all markup, styles, and JavaScript logic. This radical simplicity of deployment was a deliberate design decision: the entire application can be hosted on any static web server, distributed via email, or opened locally from a user’s filesystem. No installation, compilation, package management, or server-side processing is required. The application is currently deployed at https://descriptrongbifannotator.org via a Hetzner cloud instance (https://www.hetzner.com/) running Docker with Nginx reverse proxy and Let’s Encrypt SSL, which also serves as a CORS proxy and SAM2 encoder backend for the GBIF API.

The GBIF integration layer queries the GBIF Occurrence API v1 with pagination support, species autocompletion via the GBIF Species API, and common name resolution. Users can search by scientific name, common name, or GBIF occurrence ID, and various other criteria, with results displayed in a paginated gallery showing specimen thumbnails, scientific names, localities, and institutional codes. Upon selection, the application loads the full-resolution specimen image and automatically detects the appropriate morphological template based on taxonomic classification cascading from order through family to class level.

In addition to GBIF occurrence images, the annotator integrates with the Zenodo Biodiversity Literature Repository (BioSysLit), a curated community within Zenodo containing over 300,000 figures extracted from taxonomic publications including ZooKeys, Zootaxa, and other open-access journals (Plazi, 2026). Users can select ‘Zenodo (BioSysLit)’ from the secondary filter dropdown to search this repository by taxon name or keyword, with results filtered to image files (JPEG, PNG, TIFF). This enables annotation of published taxonomic figures, such as habitus photographs, line drawings, and SEM micrographs from species descriptions, creating a pathway to extract structured morphological data from the existing taxonomic literature. Selected images are loaded with their Zenodo DOI preserved as provenance metadata.

An additional mobile responsive design was also implemented. The interface is responsive to screen size, with a compact horizontal toolbar overlaying the canvas on devices below 900 pixels width. This mobile layout places annotation tools (point, bounding box, brush, eraser, keypoint, and segment) directly adjacent to the canvas, eliminating panel switching during annotation on tablets and phones. On small screens we find that the + and - point prompts are the best options for instance segmentation with SAM2.

Users may also optionally authenticate via manual name entry or OAuth through GBIF and iNaturalist accounts, attaching verified identities to all annotations and exports for provenance tracking.

### Morphological templates and ontology linking

A central innovation is the comprehensive library of morphological region templates organized by taxonomic group. The current release includes 25 template groups spanning all major branches of multicellular life: Formicidae, Coleoptera, Lepidoptera, Diptera, Hymenoptera, Hemiptera, Orthoptera, Odonata, Insecta (generic), Araneae, Crustacea, Myriapoda, Arthropoda (generic); Aves, Mammalia, Reptilia, Amphibia, Actinopterygii, Vertebrata (generic); Mollusca; Platyhelminthes; Angiospermae, Gymnospermae, Bryophyta, Plantae (generic); Fungi; and Lichenes. Each group provides multiple standardized views totaling 124 across the system, with every group guaranteed to include the four cardinal views: habitus lateral, habitus dorsal, habitus ventral, and head frontal (or a domain-appropriate equivalent such as cross-section for Fungi or aperture for Mollusca).

Each anatomical region within a template is linked to a formal ontology term where available, drawing from UBERON (Mungall et al., 2012) for general anatomy, the Hymenoptera Anatomy Ontology (HAO) (Yoder et al., 2010) for insect structures, the Plant Ontology (PO) (Cooper et al., 2013) for botanical features, and avian-specific terms. Regions carry associated attribute sets appropriate to their morphology, including texture, sculpture, setae density, color, color pattern, wing venation, cap shape (Fungi), thallus type (Lichenes), and 38 attribute categories each with controlled vocabularies containing 7–15 standardized values. This totals to 335 unique ontology CURIEs with 745 possible ontology instances for annotation, for a full summary see Supplementary Figures S1–S5.

### AI-assisted annotation tools

The annotation interface provides six complementary tools: (1) The system follows a split encoder/decoder design: a FastAPI backend runs the SAM2.1-Tiny encoder once per image and returns packed embeddings; the browser runs the exported ONNX mask decoder via onnx-runtime-web to update masks in response to user point prompts (positive/negative) and bounding boxes; (2) bounding box selection for coarse region identification; (3) a configurable paintbrush for manual mask creation and refinement; (4) an eraser tool for mask correction; (5) keypoint placement for landmark-based annotation; and (6) a line demarcation tool for sutures and sulci, etc. (7) A scale bar calibration tool enables users to draw a reference line along a known distance in the image, enter the real-world measurement and unit, and calibrate all subsequent measurements. The calibration (pixels per unit) is stored in all export formats, enabling downstream conversion of pixel-based measurements to physical units. (8) An edit mask mode allows users to refine existing SAM2 predictions by switching between additive painting and subtractive erasing.

The SAM2 integration uses the image encoder to generate embeddings upon image load, then performs real-time mask prediction as users interact. The lightweight decoder runs inference in under 100ms on modern hardware, enabling interactive exploration. Contour rendering uses Moore neighbourhood boundary tracing with Ramer-Douglas-Peucker simplification and Chaikin corner-cutting smoothing for publication-quality outlines. Multi-instance support allows annotating multiple specimens of the same anatomical structure (e.g., left and right antennae) with separate instance-level attributes.

**Figure 1.**
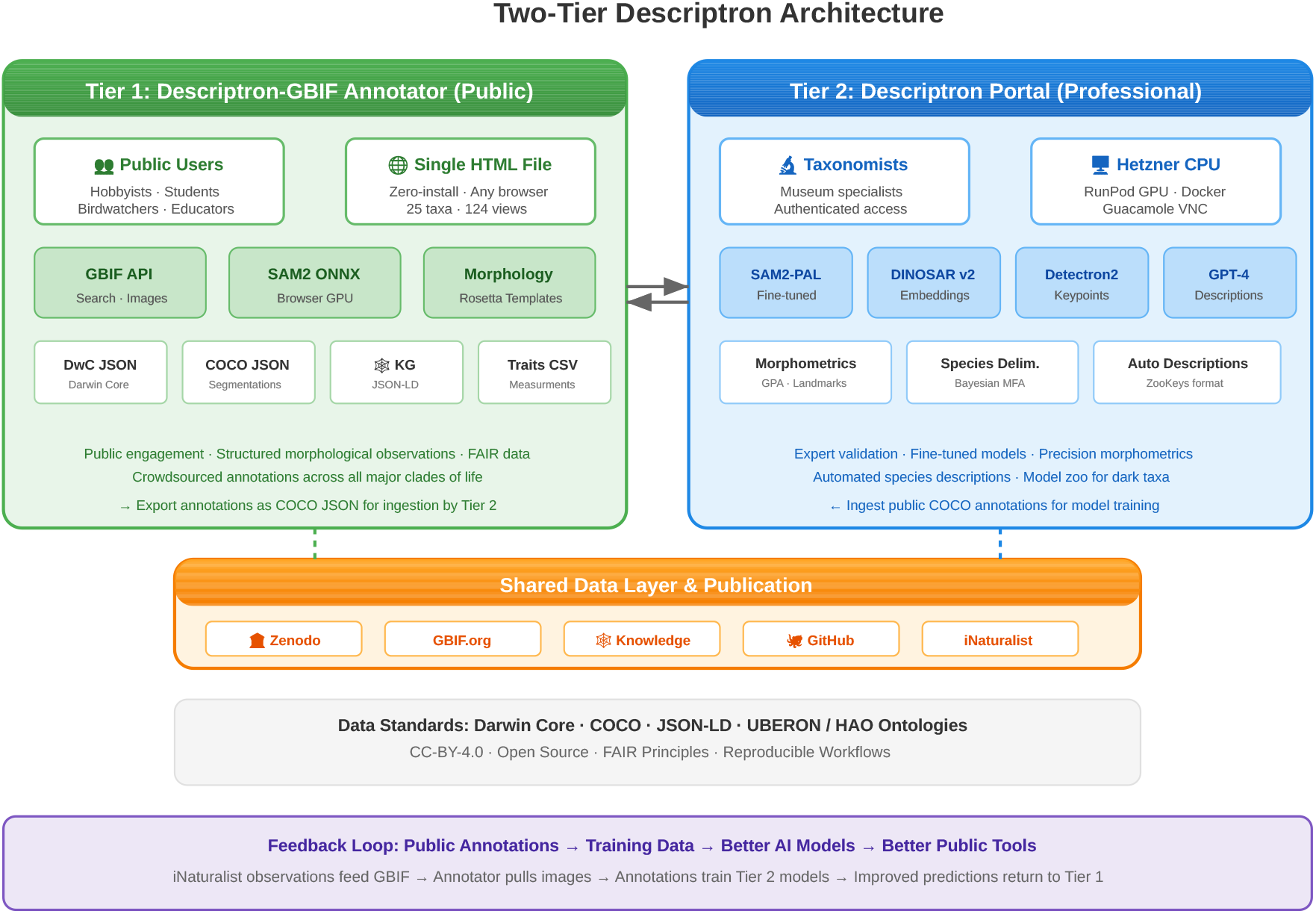
Two-tier Descriptron architecture. Tier 1 (left, green): the public-facing GBIF Annotator enables crowdsourced morphological annotation via web browser with zero installation. Tier 2 (right, blue): the professional Descriptron Portal provides GPU-accelerated tools for taxonomists including fine-tuned SAM2-PAL, DINOSARv2 embeddings, Detectron2 keypoints, and GPT-4 automated descriptions via authenticated Hetzner CPU VPS with on-demand RunPod GPU pods. Both tiers share a common data layer (orange) for publication via Zenodo, GBIF, and GitHub, with bidirectional COCO JSON exchange enabling public annotations to serve as training data for professional AI models.

### Export formats and interoperability

The annotator supports four complementary export formats designed for maximum interoperability with existing biodiversity informatics infrastructure and computer vision workflows. Darwin Core JSON packages occurrence metadata with structured measurements following the MeasurementOrFact extension (Wieczorek et al., 2012). COCO JSON exports segmentation masks, bounding boxes, keypoints, and line annotations in the standard format used by Detectron2, OpenCV and other computer vision frameworks, enabling direct use of annotations for model training (JaidedAI, 2020; Daniel Gatis and contributors, 2022; OpenCV Contributors, 2021; Wu et al., 2019). A traits CSV provides a flat tabular view of all morphological attributes suitable for statistical analysis. The knowledge graph export generates JSON-LD with a formal ontology context linking to Dublin Core, Darwin Core terms, and UBERON anatomical identifiers, producing a specimen-centric graph connecting Specimen, Taxon, Image, AnatomicalRegion, MorphologicalTrait, and SegmentationMask nodes.

Critically, the system also supports COCO JSON import, enabling bidirectional data flow with the Descriptron Portal. Annotations generated by professional taxonomists using the full desktop application can be loaded into the browser-based annotator for review, knowledge graph conversion, or Zenodo publication. The import pathway reads the template identifier from the COCO info block and loads the corresponding morphological template prior to populating annotations, ensuring correct region mapping regardless of which template is currently active.

### Zenodo integration and FAIR data

A built-in Zenodo publishing pipeline enables direct deposition of annotation datasets as citable resources with Digital Object Identifiers (DOIs). Users authenticate with a personal Zenodo access token, select which export formats to include (any combination of DwC JSON, COCO JSON, traits CSV, knowledge graph, and the specimen image itself), and can either save a draft for review or publish immediately. The deposit metadata is auto-populated from the specimen’s GBIF occurrence data including scientific name, family, institutional code, and a back-link to the GBIF occurrence as a related identifier. All deposits default to CC-BY-4.0 licensing and are tagged to the Zenodo biodiversity community. A sandbox mode enables testing without creating permanent records. This pipeline directly addresses the FAIR principles (Wilkinson et al. 2016) by making morphological annotations Findable (DOI), Accessible (open Zenodo repository), Interoperable (standard formats with ontology linking), and Reusable (CC-BY-4.0 with full provenance metadata). Each annotation session receives a unique URI (e.g., urn:descriptron:annotation:{timestamp}-{random}), and all four export formats embed both this session identifier and the originating GBIF occurrence ID or Zenodo DOI, ensuring full provenance traceability from raw image to published dataset.

**Figure 2.**
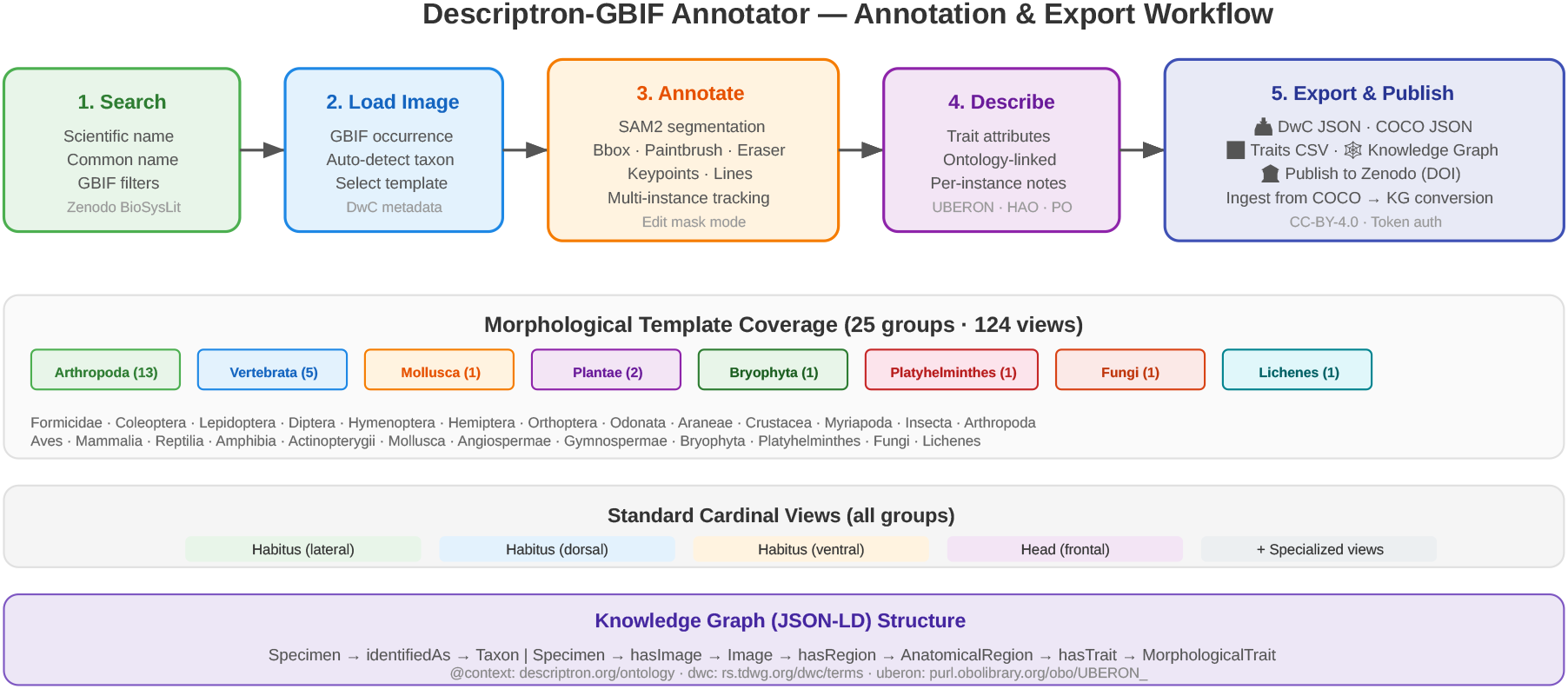
Annotation and export workflow showing the five-stage pipeline from GBIF search through structured export. The morphological template library (bottom) covers 25 taxonomic groups with 124 standardized views, each guaranteeing four cardinal perspectives. The knowledge graph structure (bottom) shows the JSON-LD entity-relationship model linking specimens to taxa, anatomical regions, and morphological traits via formal ontology identifiers.

### Two-tier architecture: from public engagement to professional taxonomy

The Descriptron-GBIF Annotator is designed as the public-facing first tier of a two-tier system. The second tier is the Descriptron Portal, a CIPRES-style science gateway (Miller et al. 2010) providing authenticated multi-user access to GPU-accelerated morphological analysis through a web browser which can be accessed here https://descriptronportal.org.

### Tier 2: The Descriptron Portal

Descriptron Portal deploys on dedicated CPU servers via Hetzner and GPU instances via RunPod (https://console.runpod.io/deploy) using Docker containerization with Apache Guacamole for browser-based VNC access, via authenticated Guacamole VNC sessions. The portal runs on a lightweight Hetzner CPU VPS with GPU compute provisioned on-demand through RunPod Secure Cloud, supporting automatic fallback across GPU types (RTX A4000 → RTX 3090 → RTX A5000 → RTX 4090 → RTX A6000) for cost-effective scaling. Each authenticated user receives an isolated container with pre-configured conda environments for Descriptron (Van Dam, 2024; Van Dam and Štarhová Serbina, 2025) which runs several different computer vision programs to help produce taxonomic species descriptions. Some environments include: SAM2-PAL (a palindrome-based mask propagation variant with cycle consistency training of SAM2 with LoRA fine-tuning), DINOSAR v2 (contrastive species-level vision embeddings), Detectron2 (instance segmentation and keypoint detection), OpenCV computer vision tools and GPT-4-API based automated description generation. A Guacamole web frontend with optional multi-factor authentication eliminates the need for VPN, SSH, or local software installation. Results are accessible via an integrated FileBrowser interface.

The Portal supports advanced workflows unavailable in the browser-based tier: fine-tuning SAM2 models on taxon-specific training data with LoRA adapters; geometric morphometrics via Generalized Procrustes Analysis with equidistant resampling and cyclic-shift optimization for rotation-invariant closed contours; multi-modal Bayesian species delimitation fusing DINOSAR v2 image embeddings, Descriptron morphological features, and DNA barcoding data with reliability gates preventing any single modality from overwhelming the fusion; and automated generation of formal species descriptions via structured GPT-4 API calls.

### The feedback loop

The two tiers are connected by a bidirectional data exchange via COCO JSON as the shared interchange format. Public annotations generated by Tier 1 users can be aggregated and ingested by the Portal for model training, progressively improving the SAM2 and Detectron2 models deployed in both tiers. Conversely, expert-validated annotations from the Portal can be exported and loaded into the public annotator for review, knowledge graph conversion, and Zenodo archival. This creates a positive feedback loop analogous to the relationship between iNaturalist community identifications and the AI models trained on them: public participation generates data that improves professional tools, which in turn improve the public experience.

## Results

### Context: citizen science and morphological data

Our approach builds on the citizen science paradigm demonstrated by Notes from Nature (Hill et al. 2012), which showed that non-specialist volunteers can make reliable contributions to biodiversity data through well-designed web interfaces with clear tasks and appropriate feedback. Where Notes from Nature focused on transcription of textual label data, the Descriptron-GBIF Annotator extends the crowdsourcing model to structured morphological observation (Fig. 3), a task that, while more complex, is facilitated by the template-driven interface, AI-assisted segmentation, and controlled vocabulary attribute systems.

**Figure 3.**
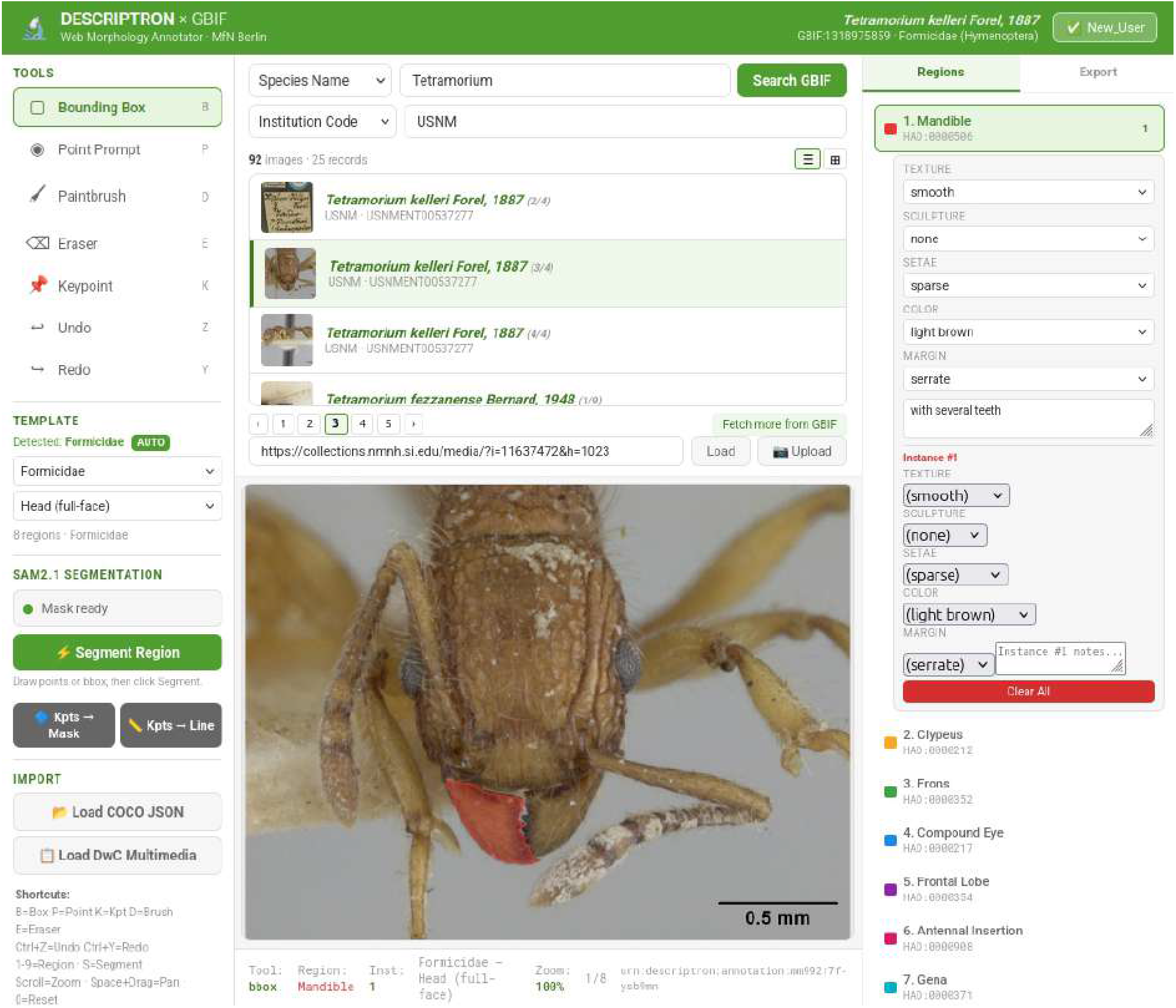
Screen-Shot of the live Descriptron-GBIF Annotater interface including a live image retrieved from the USNM of a *Tetramorium* specimen with and active annotation of the mandible including color and texture notes.

The tool is designed to serve a spectrum of users from curious hobbyists to undergraduate students in biology courses to parataxonomists and early-career researchers. For educational contexts, the structured template system functions as an interactive guide to comparative morphology: a student annotating a beetle specimen learns the terminology and spatial relationships of elytra, pronotum, head, and legs through the act of annotation itself. For research contexts, the standardized output formats and ontology linking ensure that even novice annotations can be computationally aggregated and analyzed.

The knowledge graph export is particularly significant for the emerging linked biodiversity data ecosystem. By generating JSON-LD with formal ontology references, each annotation session produces a small but well-structured fragment of a larger knowledge graph that could, in aggregate, provide the first large-scale morphological knowledge base for biodiversity, complementing the occurrence-focused data currently dominating GBIF.

### Visualizing knowledge graph exports

The JSON-LD knowledge graph files exported by the annotator can be explored using several freely available tools. The JSON-LD Playground (https://json-ld.org/playground/) validates and expands the context references. For graph visualization, Cytoscape (Shannon et al., 2003) can import the nodes and edges arrays directly for interactive network exploration. Browser-based alternatives include the d3-based JSON-LD visualizer at https://json-ld.org/visualizer/ and custom force-directed graph renderings. For SPARQL-based querying, the JSON-LD can be loaded into Apache Jena or Blazegraph for integration with broader linked data resources including Wikidata and the GBIF knowledge graph initiative. Additionally the JSON-LD could be linked to other morphology based ontology initiatives including Phenoscript (Girón et al., 2023; Montanaro and Tarasov, 2024; Tarasov et al., 2023)

## Discussion and future directions

The Descriptron-GBIF Annotator demonstrates that sophisticated morphological annotation, previously requiring desktop software and domain expertise, can be made accessible to a broad public through careful interface design, AI assistance, and template-driven workflows. Several directions for future development are planned.

First, the morphological vocabulary system will be expanded by mining published taxonomic descriptions from ZooKeys and other open-access journals using large language models and vision language models to extract character-state matrices and supplement the current template definitions. Second, we plan to develop a model zoo of pre-trained SAM2 LoRA adapters for the many of taxonomically neglected arthropod families, distributable as lightweight checkpoint files that can be loaded in the browser-based annotator for taxon-specific segmentation improvements. Third, integration with the iNaturalist API would allow direct annotation of community-uploaded photographs, extending the tool beyond museum specimens to field observations. Fourth, WebGPU-based inference could enable running larger SAM2/SAM3 model variants (Base, Large) directly in the browser for improved segmentation quality without server dependency. Fourth incorporation with other morphology based ontology programs such as Phenoscript will be essential to expand the character set available and for unifying the different ontology terms (Tarasov et al., 2023).

The two-tier architecture described here, public crowdsourcing feeder into professional AI-assisted taxonomy, represents a scalable approach to the taxonomic crisis that neither automated AI systems nor human experts alone can solve. By making morphological data collection accessible, structured, and citable, the Descriptron ecosystem aims to transform the millions of specimen images already available through GBIF from static photographs into rich, machine-readable morphological resources.

## Availability and code deposition

The Descriptron-GBIF Annotator is freely available at https://descriptrongbifannotator.org and the source code is deposited on GitHub at https://github.com/alexrvandam/Descriptron-GBIF_Annotator under an open-source license. The Descriptron Portal deployment scripts (Docker Compose, Guacamole configuration, user management) are available at https://github.com/alexrvandam/Descriptron-Portal and with the main portal accessible here: https://descriptronportal.org. The complete codebase including the HTML annotator, CORS proxy server, and deployment documentation will archived on Zenodo via GitHub.

## Acknowledgments

We thank the Museum für Naturkunde Berlin and the Center for Integrative Biodiversity Discovery for institutional support. We acknowledge the GBIF Secretariat for maintaining the data infrastructure that makes this tool possible, and the Meta AI Research team for the open-source release of SAM2. We thank the referees for their useful comments and time taken to review this work.

## Supplementary Figures

**Supplementary Figure S1.**
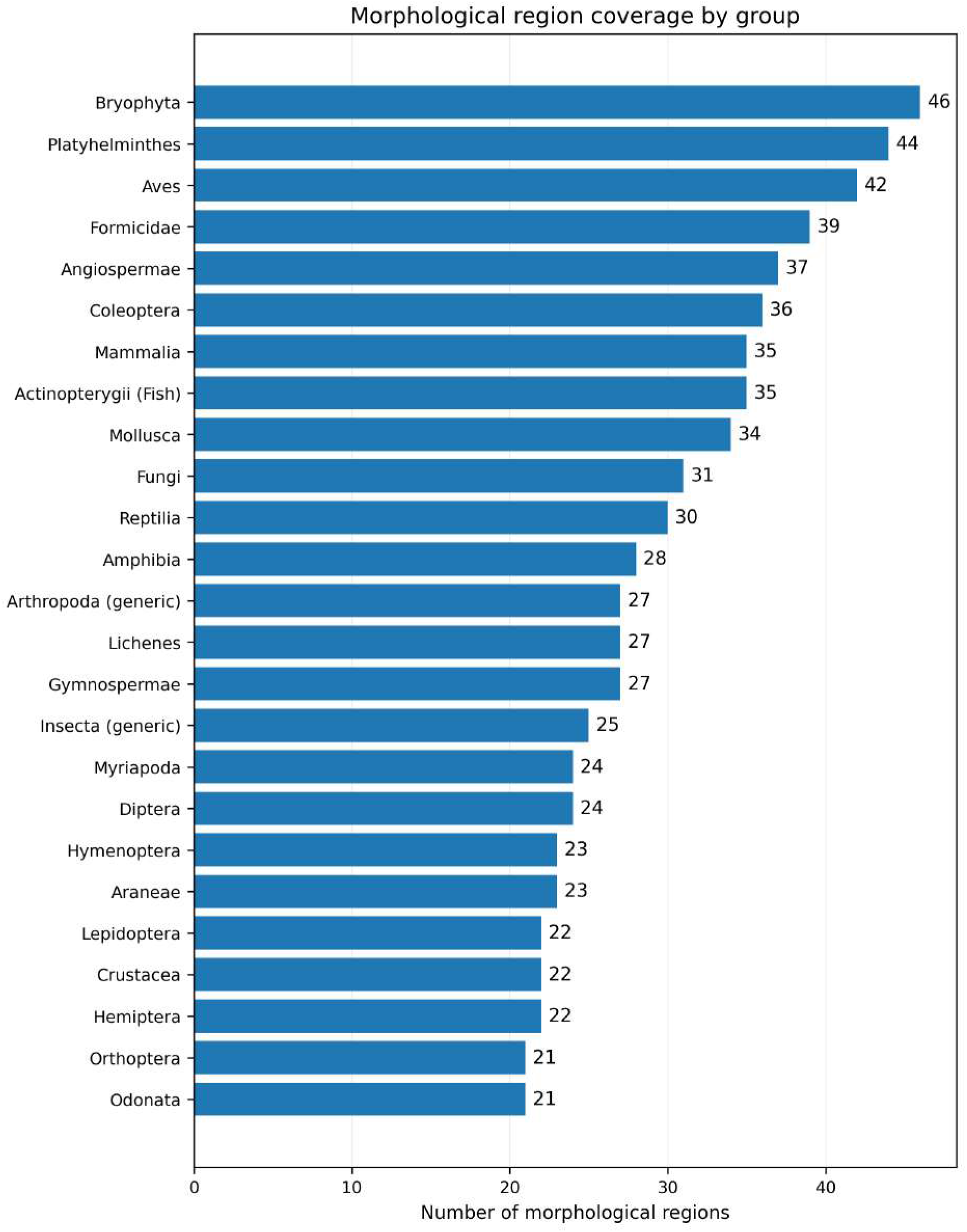
“Morphological region coverage by group”. Horizontal bar chart showing the number of morphological regions defined/covered for each taxonomic group (x-axis: number of morphological regions; y-axis: group). Values are annotated at the end of each bar.

**Supplementary Figure S2.**
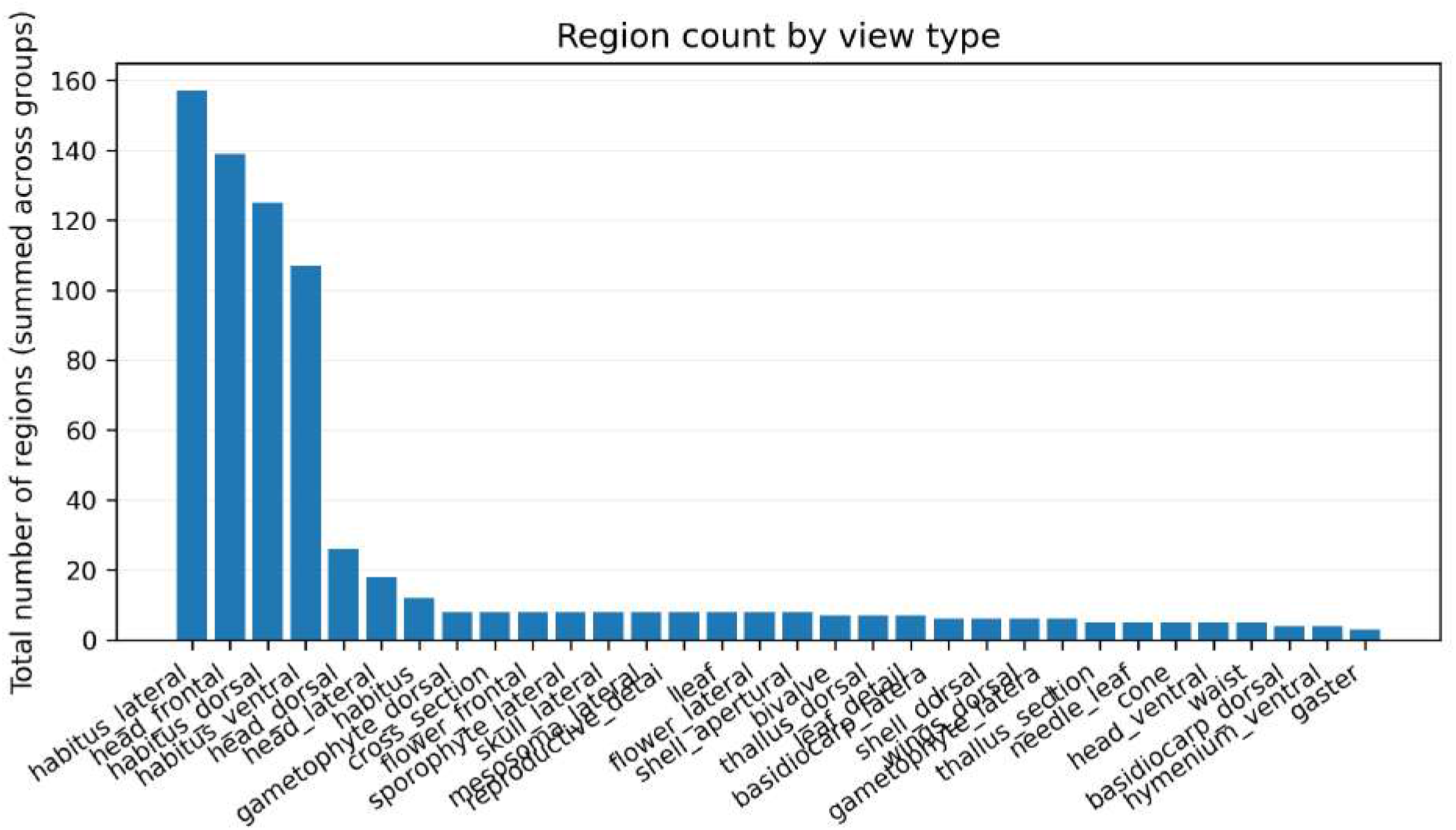
“Region count by view type”. Bar chart summarizing the total number of regions (summed across groups) for each view type (x-axis: view type; y-axis: total number of regions). View types are ordered from highest to lowest total.

**Supplementary Figure S3.**
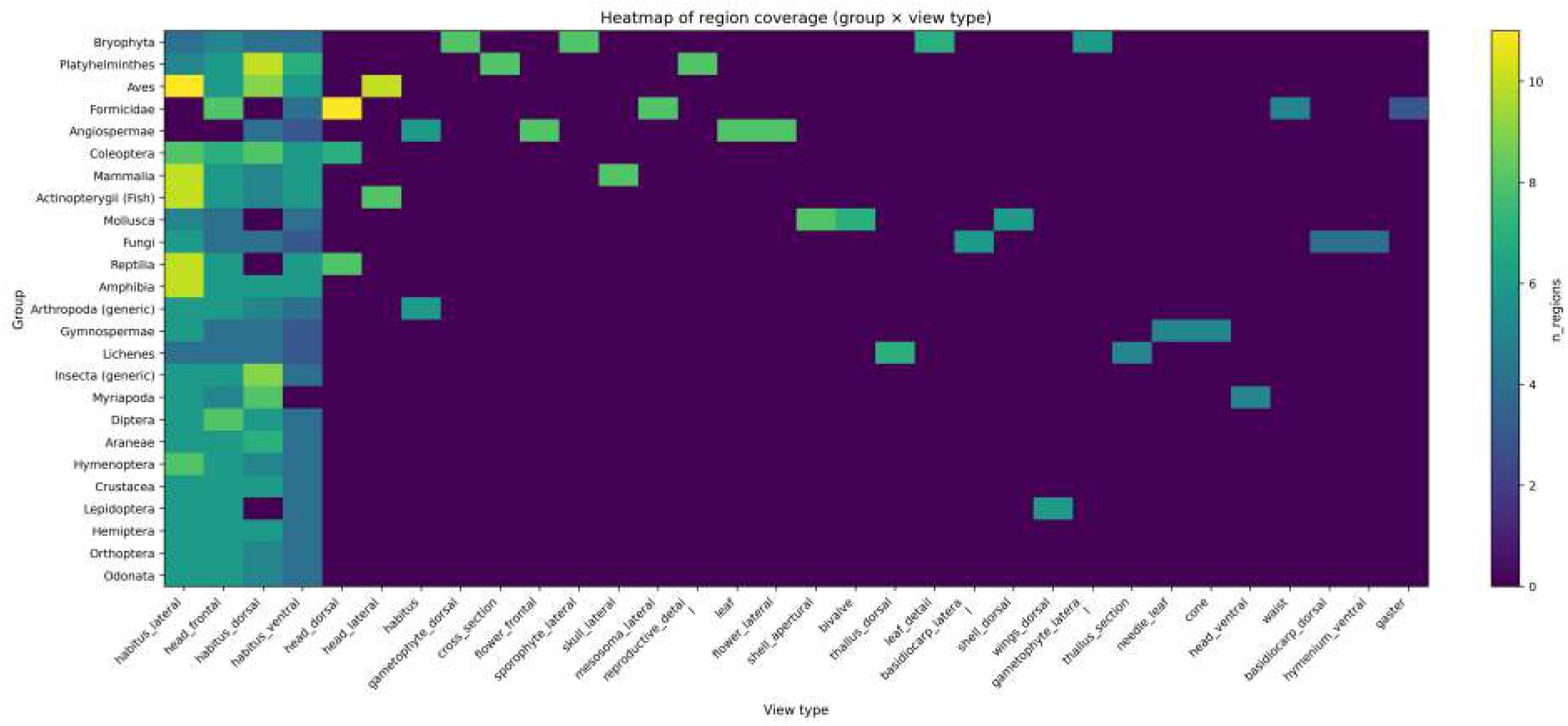
“Heatmap of region coverage (group × view type)”. Heatmap showing n_regions for each group × view type combination (x-axis: view type; y-axis: group). Cell color encodes the number of regions (colorbar labeled n_regions).

**Supplementary Figure S4.**
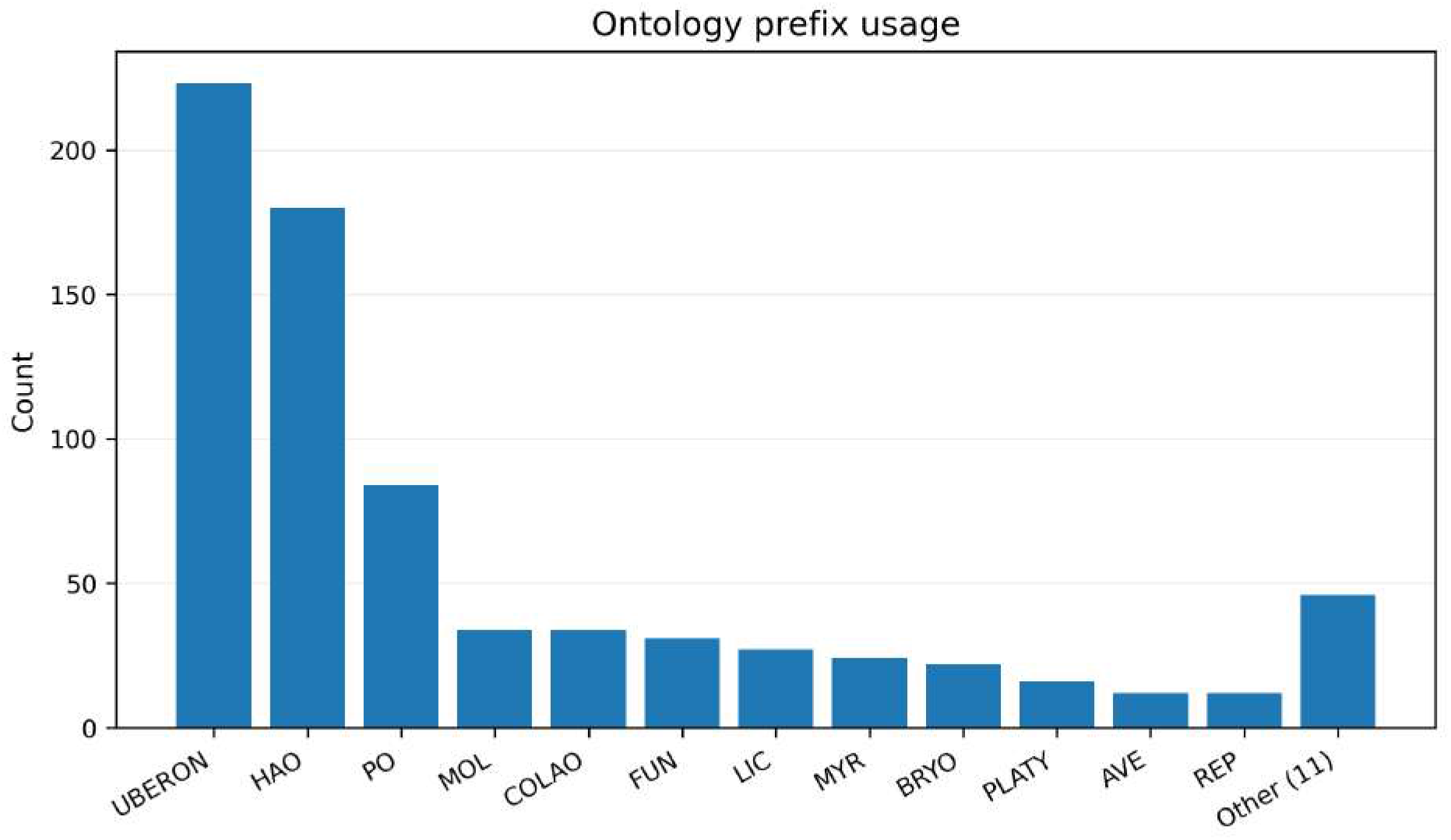
“Ontology prefix usage”. Bar chart of the count of ontology CURIE prefixes used (e.g., UBERON, HAO, PO, etc.), with “Other (11)” aggregating prefixes outside the main set (x-axis: prefix; y-axis: count).

**Supplementary Figure S5.**
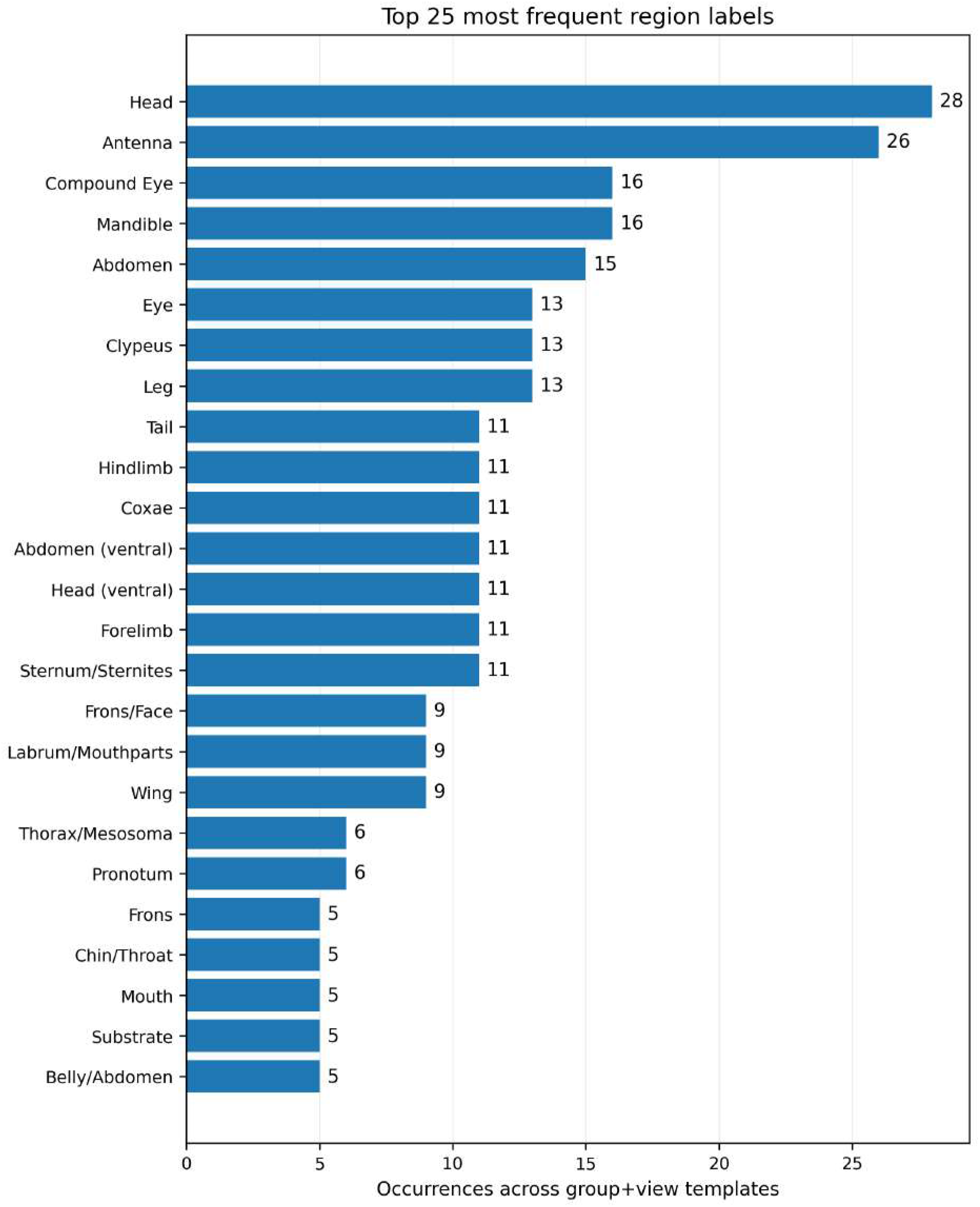
“Top 25 most frequent region labels”. Horizontal bar chart of the 25 most frequent region labels and their occurrence counts across group+view templates (x-axis: occurrences; y-axis: region label). Values are annotated at the end of each bar.

## Notes

https://descriptrongbifannotator.org

https://descriptronportal.org

https://github.com/alexrvandam/Descriptron-GBIF_Annotator

https://github.com/alexrvandam/Descriptron-Portal

### Competing Interest Statement

The authors have declared no competing interest.

## References

Cooper, L., Walls, R.L., Elser, J., Gandolfo, M.A., others, 2013. The Plant Ontology as a tool for comparative plant anatomy and genomic analyses. Plant & Cell Physiology 54, e1. 10.1093/pcp/pcs163

Daniel Gatis and contributors, 2022. rembg: Image background removal.

Engel, M.S., Ceríaco, L.M.P., Daniel, G.M., Dellapé, P.M., Diller, J., Jr., C.E., others, 2021. The taxonomic impediment: a shortage of taxonomists, not of taxonomy. Zoological Journal of the Linnean Society 193, 381–447. 10.1093/zoolinnean/zlaa061

GBIF, 2026. Global Biodiversity Information Facility. GBIF.org [Data portal]. Retrieved March 10, 2026, from GBIF.org

GBIF, 2011. GBIF: The Global Biodiversity Information Facility (2011) What is GBIF?. [WWW Document]. URL Available from https://www.gbif.org/what-is-gbif [13 January 2020].

Girón, J.C., Tarasov, S., González Montaña, L.A., Matentzoglu, N., Smith, A.D., Koch, M., Boudinot, B.E., Bouchard, P., Burks, R., Vogt, L., Yoder, M., Osumi-Sutherland, D., Friedrich, F., Beutel, R.G., Mikó, I., 2023. Formalizing Invertebrate Morphological Data: A Descriptive Model for Cuticle-Based Skeleto-Muscular Systems, an Ontology for Insect Anatomy, and their Potential Applications in Biodiversity Research and Informatics. Systematic Biology 72, 1084–1100. 10.1093/sysbio/syad025

Hartop, E., Srivathsan, A., Ronquist, F., Meier, R., 2022. Towards Large-Scale Integrative Taxonomy (LIT): Resolving the Data Conundrum for Dark Taxa. Systematic Biology 71, 1404–1422. 10.1093/sysbio/syac033

Hill, A., Guralnick, R., Smith, A., Sallans, A., Gillespie, R., Denslow, M., Gross, J., Murrell, Z., Conyers, T., Oboyski, P., Ball, J., Thomer, A., Prys-Jones, R., de la Torre, J., Kociolek, P., Fortson, L., 2012. The notes from nature tool for unlocking biodiversity records from museum records through citizen science. ZooKeys 209, 219–233. 10.3897/zookeys.209.3472

iNaturalist, 2026. iNaturalist. Available from https://www.inaturalist.org.Accessed 10 March x2026.10.15468/ab3s5x

JaidedAI, 2020. EasyOCR: Ready-to-use OCR with 80+ supported languages. https://github.com/JaidedAI/EasyOCR

Miller, M.A., Pfeiffer, W., Schwartz, T., 2010. Creating the CIPRES Science Gateway for inference of large phylogenetic trees, in: 2010 Gateway Computing Environments Workshop (GCE). IEEE, pp. 1–8. 10.1109/GCE.2010.5676129

Montanaro, G., Balhoff, J.P., Girón, J.C., Söderholm, M., Tarasov, S., 2024. Computable species descriptions and nanopublications: applying ontology-based technologies to dung beetles (Coleoptera, Scarabaeinae). Biodiversity Data Journal 12, e121562. 10.3897/BDJ.12.e121562

Mora, C., Tittensor, D.P., Adl, S., Simpson, A.G.B., Worm, B., 2011. How Many Species Are There on Earth and in the Ocean? PLOS Biology 9, 1–8. 10.1371/journal.pbio.1001127

Mungall, C.J., Torniai, C., Gkoutos, G.V., Lewis, S.E., Haendel, M.A., 2012. Uberon, an integrative multi-species anatomy ontology. Genome Biology 13, R5. 10.1186/gb-2012-13-1-r5

OpenCV Contributors, 2021. OpenCV: Open Source Computer Vision Library.

Oquab, M., Darcet, T., Moutakanni, T., Vo, H., Szafraniec, M., Khalidov, V., Fernandez, P., Haziza, D., Massa, F., El-Nouby, A., others, 2023. DINOv2: Learning Robust Visual Features without Supervision. arXiv preprint.

Plazi, 2026. Biodiversity Literature Repository. Zenodo. https://zenodo.org/communities/biosyslit/

Ravi, N., Gabeur, V., Hu, Y.-T., Hu, R., Ryali, C., Ma, T., Khedr, H., Rädle, R., Rolber, C. Gustafson, L., Mintun, E., Pan, J., Alwala, K.V., Carion, N., Wu, C.-Y., Girshick, R., Dollár, P., Feichtenhofer, C., 2024. SAM 2: Segment Anything in Images and Videos. arXiv preprint arXiv:2408.00714. 10.48550/arXiv.2408.00714

Shannon, P., Markiel, A., Ozier, O., Baliga, N.S., Wang, J.T., Ramage, D., Amin, N., Schwikowski, B., Ideker, T., 2003. Cytoscape: a software environment for integrated models of biomolecular interaction networks. Genome Research 13, 2498–2504. 10.1101/gr.1239303

Siméoni, O., Vo, H.V., Seitzer, M., Baldassarre, F., Oquab, M., Jose, C., Khalidov, V., Szafraniec, M., Yi, S., Ramamonjisoa, M., Massa, F., Haziza, D., Wehrstedt, L., Wang, J., Darcet, T., Moutakanni, T., Sentana, L., Roberts, C., Vedaldi, A., Tolan, J., Brandt, J., Couprie, C., Mairal, J., Jégou, H., Labatut, P., Bojanowski, P., 2025. DINOv3. arXiv preprint arXiv:2508.10104.

Tarasov, S., 2023. PhenoScript [Software]. GitHub. https://github.com/sergeitarasov/PhenoScript

Van Dam, A.R., 2024. Descriptron [WWW Document]. URL https://github.com/alexrvandam/Descriptron

Van Dam, A.R., Štarhová Serbina, L., 2025. Descriptron: Artificial intelligence for automating taxonomic species descriptions with a user-friendly software package. Systematic Entomology e70005. 10.1111/syen.70005

Van Horn, G., Mac Aodha, O., Song, Y., Cui, Y., Sun, C., Shepard, A., Adam, H., Perona, P., Belongie, S., 2018. The iNaturalist Species Classification and Detection Dataset, in: 2018 IEEE/CVF Conference on Computer Vision and Pattern Recognition. Presented at the 2018 IEEE/CVF Conference on Computer Vision and Pattern Recognition (CVPR), IEEE, Salt Lake City, UT, pp. 8769–8778. 10.1109/CVPR.2018.00914

Wheeler, Q.D., Raven, P.H., Wilson, E.O., 2004. Taxonomy: Impediment or Expedient? Science 303, 285. 10.1126/science.1092363

Wieczorek, J., Bloom, D., Guralnick, R., Blum, S., Döring, M., Giovanni, R., Robertson, T., Vieglais, D., 2012. Darwin Core: An Evolving Community-Developed Biodiversity Data Standard. PLoS ONE 7, e29715. 10.1371/journal.pone.0029715

Wilkinson, M.D., Dumontier, M., Aalbersberg, I.J., Appleton, G., Axton, M., Baak, A., others, 2016. The FAIR Guiding Principles for scientific data management and stewardship. Scientific Data 3, 160018. 10.1038/sdata.2016.18

Wu, Y., Kirillov, A., Massa, F., Lo, W.-Y., Girshick, R., 2019. Detectron2. https://github.com/facebookresearch/detectron2

Yoder, M.J., Mikó, I., Seltmann, K.C., Bertone, M.A., Deans, A.R., 2010. A gross anatomy ontology for Hymenoptera. PLoS ONE 5, e15991. 10.1371/journal.pone.0015991

